# Multiplane HiLo microscopy with speckle illumination and non-local means denoising

**DOI:** 10.1101/2023.09.08.556851

**Authors:** Shuqi Zheng, Minoru Koyama, Jerome Mertz

## Abstract

**Significance:** HiLo microscopy synthesizes an optically-sectioned image from two images, one obtained with uniform and another with patterned illumination, such as laser speckle. Speckle-based HiLo has the advantage of being robust to aberrations, but is susceptible to residual speckle noise that is difficult to control. We present a computational method to reduce this residual noise without compromising spatial resolution. In addition, we improve the versatility of HiLo microscopy by enabling simultaneous multiplane imaging (here 9 planes).

**Aim:** Our goal is to perform fast, high contrast multiplane imaging with a conventional camera-based fluorescence microscope.

**Approach:** Multiplane HiLo imaging is achieved with the use of a single camera and z-splitter prism. Speckle noise reduction is based on the application of a non-local means (NLM) denoising method to perform ensemble averaging of speckle grains.

**Results:** We demonstrate the capabilities of multiplane HiLo with NLM denoising both with synthesized data and by imaging cardiac and brain activity in zebrafish larvae at 40 Hz frame rates.

**Conclusions:** Multiplane HiLo microscopy aided by NLM denoising provides a simple tool for fast, optically-sectioned volumetric imaging that can be of general utility for fluorescence imaging applications.

## 1 Introduction

Camera-based widefield microscopy enables fluorescence imaging with larger field of view, higher speed and higher sensitivity than standard scanning-based methods. However, in its most simple configuration, camera-based widefield microscopy fails to provide optical sectioning, which undermines imaging contrast particularly for thick samples.

Widefield optical sectioning techniques have been developed by engineering either the illumination^1^ or detection^2^ of a microscope, or both.^3, 4^ A commonly adopted technique is structured illumination microscopy (SIM),^5^where a sequence of grid patterns (at least three) is projected into the sample and the emitted fluorescence is imaged by a camera. The modulations of the grid pattern are resolvable only when they are in focus, meaning that optical sectioning can be obtained by simple demodulation.^5, 6^

HiLo microscopy^7–9^is a variant of SIM. Instead of ensuring full illumination coverage from a sequence of structured illumination patterns, HiLo directly makes use of a uniform-illumination image, to which it adds sectioning information obtained from a single additional structured illumination (with either a grid^9^ or speckle pattern^7, 8^), enabling optical sectioning with only two shots. Speckle has an advantage of providing intrinsically high contrast, with intensity standard deviations as large as the average intensity itself. Moreover, HiLo with speckle illumination is simpler to implement and much more resistant to aberrations than grid illumination. However, speckle is highly noisy, with substantial noise residing not only in the intensity but also in the variance of the intensity, from which the sectioning information is extracted. This higher-order noise is difficult to control, and in our original HiLo algorithm produced considerable residual noise artifacts in our final processed images. The simplest way to mitigate this residual noise is with spatial filtering, however this comes at the expense of reduced optical sectioning capacity.^10^ Alternatively, one could consider using speckle with customized intensity statistics.^11^ Here, to mitigate the tradeoff between image fidelity and sectioning strength, we propose a computational method based on a non-local means (NLM) denoising algorithm^12^ to reduce the speckle noise in our final HiLo reconstruction.

Denoising by NLM is different from local smoothing. Instead, it takes advantage of image redundancy and seeks out similar small windows, or patches, (not necessarily local) with different noise realizations, which can be matched and processed together through averaging. In practice, patch similarities are determined and then mapped into weights to effectively perform a weighted average of noisy pixels, providing a more robust estimate of a denoised image. NLM denoising has been applied to speckle reduction in coherent imaging systems,^13–16^ where patches of the same underlying object but modulated by different speckle realizations are ensemble averaged. Somewhat more involved, transforms can be applied to the multiplicative speckle noise in order to adapt it to additive Gaussian-noise-based NLM for when determining patch similarity.^17^ Here, rather than seeking out non-local patch similarities within our speckle-illumination image, we seek them out within our uniform illumination image, rendering the similarity search more robust and enabling the use of an already well established denoising algorithm.^18^ The similarity weights obtained from our uniform illumination image are then applied to both the high and low frequency components in our HiLo processing algorithm, thus leading to reduced noise artifacts in our final HiLo reconstruction.

Another challenge in fluorescence microscopy is volumetric imaging. In conventional scanning- and camera-based methods, axial scanning is generally required to perform imaging over extended depth ranges, thus limiting acquisition speed. To address this challenge, multifocus imaging methods have been developed that simultaneously project images from multiple depths onto a single camera with the use of an engineered grating,^19^ or cascaded prisms and beam splitters,^20, 21^ though without the benefit of background rejection. Variants incorporating optical sectioning have made use of structured illumination^22, 23^ or light sheet illumination in multi-slit^24, 25^ or multiplane^26^ geometry.

Here, we find that optically-sectioned multiplane imaging can naturally be performed by HiLo. That is, in addition to conferring NLM-based denoising to our speckle-based HiLo algorithm, we enable the capacity of multiplane HiLo imaging by the use of a z-splitter prism.^21^ We demon-strate the performance of our system first with numerical simulations and then with both fixed and live samples. In particular, we perform simultaneous optically-sectioned multiplane fluorescence imaging both of fast cardiac motion and of brain activity in larval zebrafish.

## 2 HiLo reconstruction and denoising

### 2.1 Basic principle of HiLo

In our original speckle-based HiLo algorithm,^8^two raw images *I*_*s*_(***ρ***) and *I*_*u*_(***ρ***) are acquired with speckle and uniform illumination respectively (***ρ*** is the 2D spatial coordinate at the camera plane). A final optically-sectioned image is synthesized from the fusion of the two, where sectioning of low spatial frequency components is based on the contrast decay of the imaged speckle as a function of defocus. This contrast decay can be accelerated with the help of a wavelet-type filter.^8, 27^ Specifically, contrast can be defined as

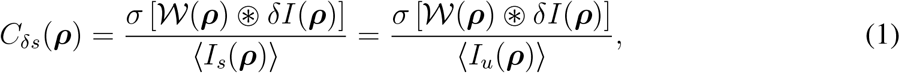

where *𝒲* (***ρ***) is a wavelet filter, δI(***ρ***) = *I*_*s*_(***ρ***)*−I*_*u*_(***ρ***) is a difference image, ⊛ denotes convolution and σ[*·*] denotes standard deviation (see Material and methods).

*C*_*δs*_(***ρ***) serves as a weighting function that preferentially extracts the optically-sectioned infocus contributions in *I*_*u*_(***ρ***). An estimate of these contributions is given by *I*_*cu*_(***ρ***) = *C*_*δs*_(***ρ***)*I*_*u*_(***ρ***), however this estimate is noisy due to residual speckle noise. The simplest strategy to control this noise is by the use of a low-pass filter, leading to a smoothed estimate of the in-focus contributions for low spatial frequencies, given by *I*_Lo_(***ρ***) = LP [*I*_*cu*_(***ρ***)]. The complementary high-frequency components, which are inherently optically-sectioned, are obtained by high-pass filtering the uni-form image: *I*_Hi_ = HP [*I*_*u*_], where 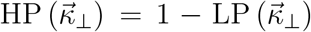. The final HiLo image, optically sectioned over the full spatial frequency bandwidth, is synthesized by combining Hi and Lo components with a scaling factor η

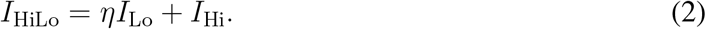

### 2.2 HiLo with non-local means denoising

#### 2.2.1 *I*_Lo_ *with NLM denoising*

The sectioning capacity of HiLo can be tuned by the cutoff frequency separating Hi and Lo. The larger the frequency, the tighter the optical sectioning, but also the more residual speckle noise becomes apparent in *I*_Lo_ because it is only weakly filtered from *I*_*cu*_ (see Fig 1(a)). To aid this filtering, we incorporate the method of NLM denoising.^12^ The basic idea is to expand the regions over which filtering is performed without lowering the cutoff frequency. This is achieved by searching for similar patches throughout *I*_*cu*_ (guided here by the uniform image) and performing a weighted average of these according to their similarity. The net result is that the residual speckle noise is substantially decreased while maintaining tight optical sectioning. Figure 1(a) illustrates this process using a numerically synthesized test target, where the out-of-focus background is simulated from the same target rotated by 180 degrees. The algorithm takes two inputs, one *I*_*u*_ from uniform illumination and the other *I*_*cu*_ from the pre-processed difference image according to Eq. 1.

**Fig 1.**
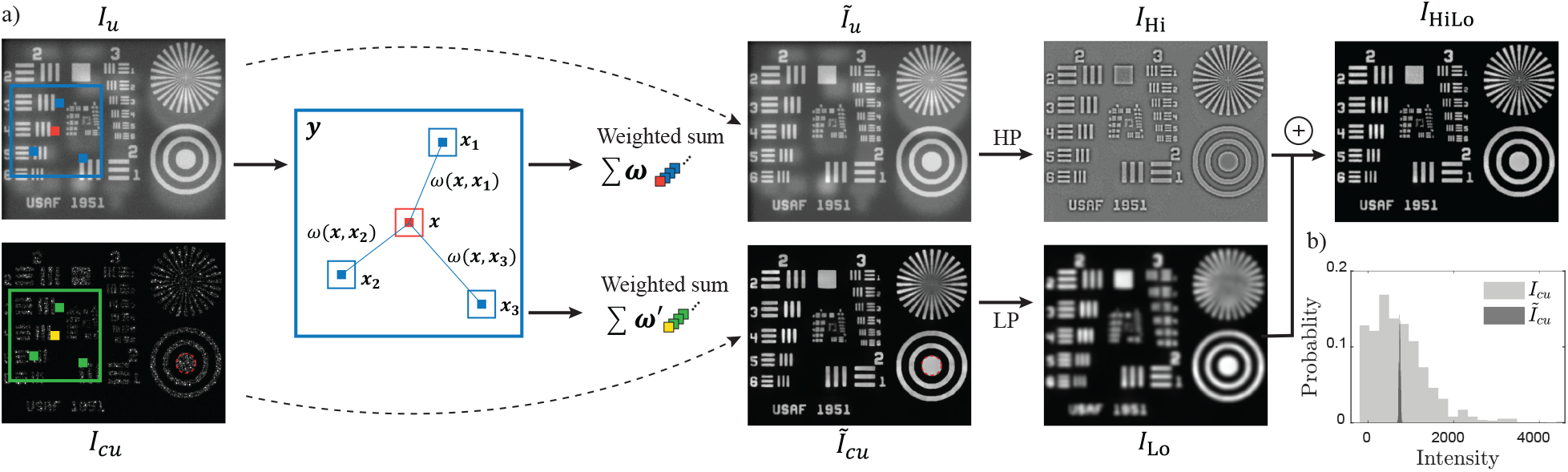
Diagram of NLM-assisted HiLo. (a) For input images *I*_*u*_ and *I*_*cu*_, weighted averagings of similar non-local patches obtained from *I*_*u*_ produce denoised 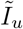 and 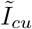. The intermediate results are then high-pass/low-pass filtered respectively to form *I*_Hi_, *I*_Lo_, which are then fused to synthesize a final optically-sectioned HiLo image *I*_HiLo_. (b) Comparison of pixel intensity distributions in the circled regions in *I*_*cu*_ and 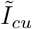.

First, similar patches are found in the uniform image *I*_*u*_. The notion of similarity comes from treating a noisy patch as a realization of a random variable of given distribution (e.g. Poisson distribution in the case of shot noise), where the parameters of the distribution are determined by the underlying noise-free patch.A pair of noisy patches are considered similar when they obey the same distribution of common parameters.^12^ Consider a vectorized square patch ***x*** containing (2N + 1)^2^ pixels centered on pixel *x*, where the center pixel has value *I*_*u*_(*x*) and the remaining pixels have value *I*_*u*_(*x* + d*x*), with dx being the pixel index across the patch ***x*** (d*x ∈* [*−*2N(N + 1), 2N(N + 1)]). For a pair of patches (***x***_**1**_, ***x***_**2**_) in the uniform image *I*_*u*_ corrupted by Poisson noise, the patch similarity is calculated based on the log likelihood

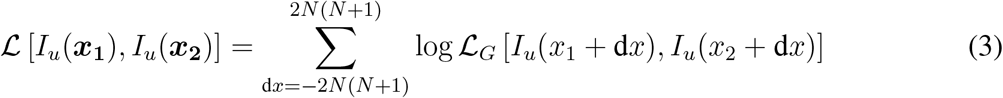

where *ℒ*_*G*_ is a pixel-wise implementation of the generalized likelihood ratio (GLR),^28^ which has been demonstrated to provide the best estimate in the case of strong noise levels and reasonably good estimates in the case of medium/low noise levels as compared to other metrics such as Euclidean distance.^28^

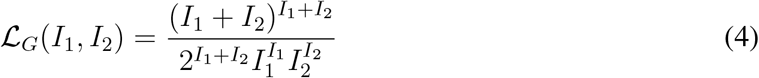

*ℒ*_*G*_ equals 1 for identical pixel values and varies between (0, 1) for different pixels.

Following Eq. 3, the weights are calculated by introducing a filtering parameter *h* that controls the degree of averaging between non-local patches, obtaining

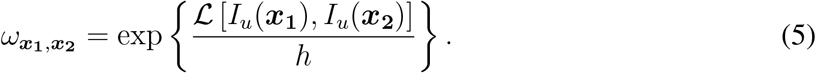

For a pixel *x*_1_ to be estimated, 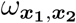 determines the contribution of pixel *x*_2_ in the weighted average. The number of patches used for calculation is further defined by a large search window ***y*** containing (2*N*_*y*_ + 1)^2^ pixels (blue rectangle in Fig. 1(a)).

Finally, the same pixel location is found in *I*_*cu*_ and the weighted average of *I*_*cu*_ is performed for all pixels within ***y***:

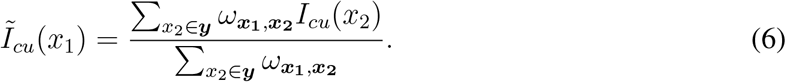

We observe that Eq. 6 is essentially an incoherent averaging of speckle realizations in *I*_*cu*_, taking into account of sample variation, which is a common practice for suppressing speckle.^29^ Figure 1(b) compares the intensity distribution in *I*_*cu*_ and 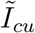 before and after the application of NLM denoising in the circled region in Fig. 1(a) where the intensity is uniform. NLM denoising significantly reduces the standard deviation of the residual speckle contrast, thus improving the local smoothness of *I*_*Lo*_.

#### 2.2.2 I_Hi_ with NLM denoising

As noted above, tighter optical sectioning is obtained with higher cutoff frequency between Hi and Lo. *I*n addition to exacerbating the problem of residual noise in *I*_Lo_, a high cutoff frequency also leads to reduced signal to noise ratio (SNR) in *I*_Hi_. Here, too, NLM denoising can be used to advantage, with little added computational burden. Specifically, the same weights are calculated from Eq. 5 with a second weighting parameter *h^′^* and the weighted averages of patches from *I*_*u*_ are applied to obtain 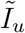, which is a denoised version of the uniform image.

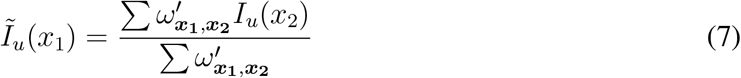

The denoised high-frequency component is then obtained by 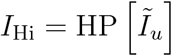.

## 3 Materials and methods

### 3.1 Multiplane imaging setup

A schematic of the experimental setup is shown in Fig. 2. HiLo requires fast switching between speckle and uniform illumination in an otherwise conventional widefield microscope. This was obtained here with two separated beams combined by a polarizing beam splitter (PBS). The first beam was produced by a collimated blue LED (SOLIS-470C); the second beam was from a continuous wave laser (Vortran Stradus, 488nm) incident on a static diffuser conjugate to the back pupil of the microscope objective (Olympus UMPLFLN 20XW or LUMPLFLN 40XW). Both light sources were triggered by a NI-DAQ (National Instruments, NI USB-6356) to produce alternating illumination synchronized to the camera exposure (PCO.edge 4.2). We note that in previous single-beam designs, the uniform illumination image was obtained by rapidly randomizing the speckle illumination patterns with a tilting or rotating diffuser.^7, 30^ Our current two-beam design avoids two potential drawbacks. First, the toggling between speckle and uniform illumination is no longer slowed down by any need to average over multiple speckle patterns. Second, the speckle illumination pattern in our case remains fixed from frame to frame, meaning that it does not introduce temporal fluctuations from frame to frame that could perturb measurements of *relative* changes in fluorescence, as utilized, for example, with calcium imaging. Finally, to perform simultaneous multiplane imaging, the emitted fluorescence was directed into a 9-plane z-splitter prism^21^ and projected onto the single camera sensor with a relay lens pair (demagnification 250/90).

**Fig 2.**
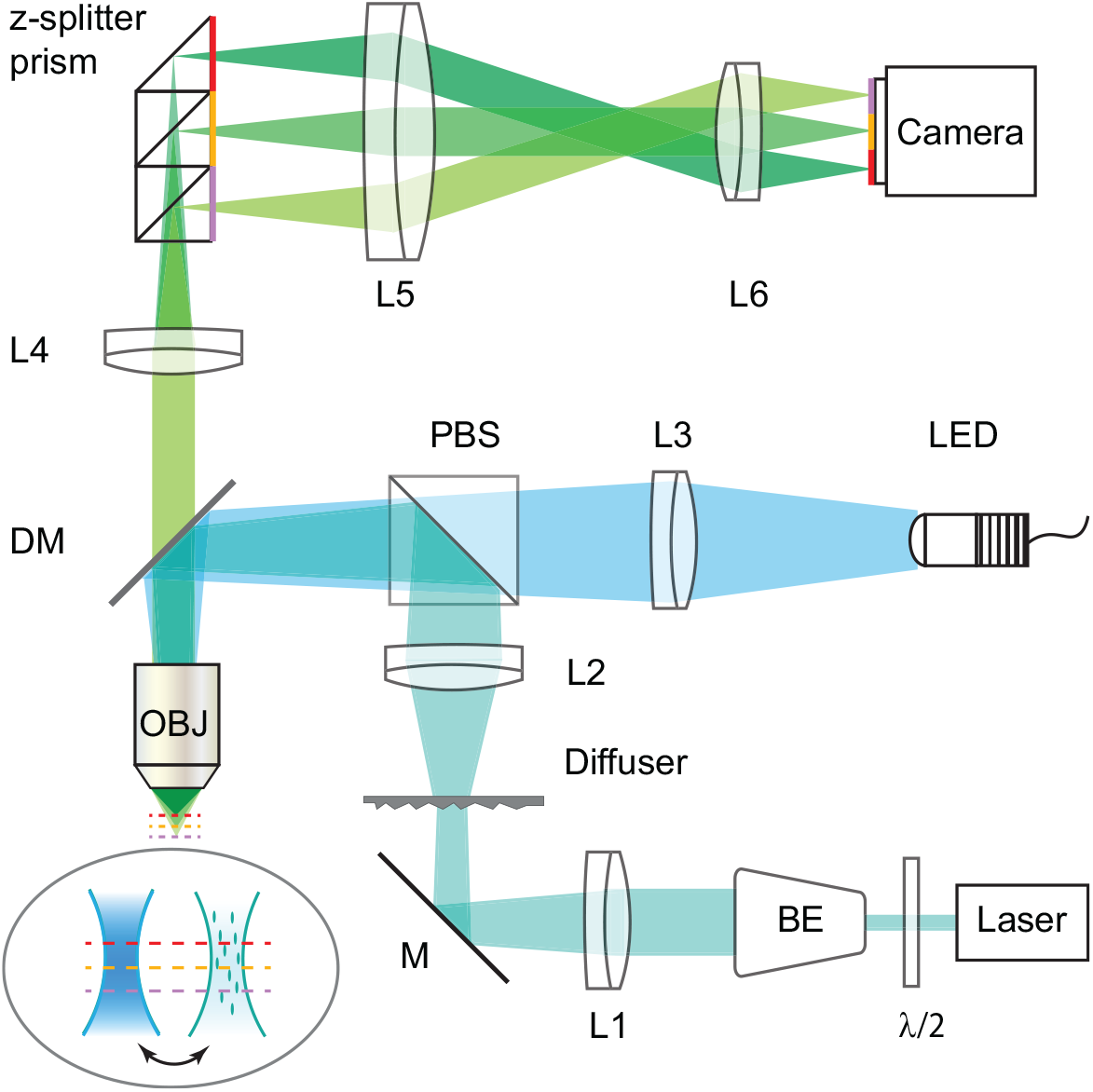
Experimental setup. Light generated from a LED and laser are recombined with a polarizing beamsplitter, providing uniform and speckle illumination of the sample. A z-splitter prism (here, shown as a 3-plane prism) enables simultaneously imaging of multiple depths with a single camera.

### 3.2 Sample preparation

*In-vivo* imaging was performed of both the heart (isl2b:Gal4 UAS:Dendra, 8dpf) and brain (elavl3:H2BGCaMP6f, 6dpf) of transgenic zebrafish larvae. To prepare for imaging, larvae were embedded in 1.5% low melting point agarose gelled on a petri dish. Larvae were positioned right-side down for heart imaging and ventral-side down for brain imaging. The solidified agarose and fish were immersed in fish water before being transferred onto the sample stage for imaging.

### 3.3 Image processing

The wavelet filter is defined in frequency space as

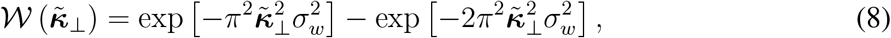

where 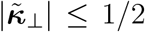 is the normalized spatial frequency. Equation 8 corresponds to *𝒲*(***ρ***) in the spatial domain

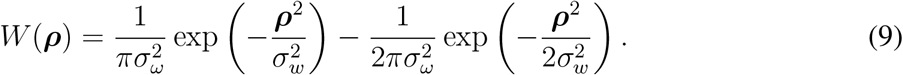

The default value of σ_*w*_ was set such that the width of the wavelet filter was about one speckle-grain size. In the case of low signal level, the wavelet size was set to slightly larger than the speckle-grain size to improve SNR. Bias introduced by shot noise and readout noise was subtracted from the measured variance,^8^ presuming a camera gain of 0.46 e^*−*^/ADU and readout noise of 1.2 e^*−*^. The scaling factor η was determined either theoretically from system parameters^8^ or empirically based on intensity distributions.

For the implementation of NLM, the patch size was chosen to be 3*×*3 pixels in all experiments, corresponding to about one speckle grain per patch, and the search window size was chosen in the range 15 *×* 15 to 61 *×* 61 pixels depending on the speckle contrast. A larger search window allows the averaging of more patches (i.e. more uncorrelated speckle) but also takes longer to process. The filtering parameters *h* for *I*_*Lo*_ and *h*^*′*^ for *I*_*Hi*_ were empirically set to be around 5 to 20 and 0.4 to 0.6 to make allowances for different speckle and noise variances. The NLM implementation was performed with a compiled C^++^ mex-function in parallel. Other pre/post-processing was performed using MATLAB 2021a.

To characterize the quality of HiLo reconstructions, we made use of the peak signal to noise ratio (PSNR), defined as

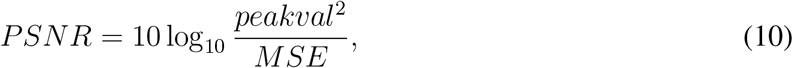

where *MSE* is the mean square error between test and reference images.

## 4 Results

### 4.1 Simulation results

We first simulated widefield imaging of a pollen grain under uniform and speckle illumination. The ground truth was obtained from a confocal image stack of an actual pollen grain obtained with Olympus FV3000 (60*×*, 1.2NA). To simulate widefield images, the volume (323*×*323*×*35 pixels, 42*×*42*×*19.6 μm^3^) was convolved with a simulated 3D point spread function (PSF) and the defocused planes produced the background for each focused slice. Shot noise and readout noise were introduced assuming a camera gain of 2.5 [ADU/e^*−*^] and readout noise of 1.6 e^*−*^ (rms).

Figure 3 shows the simulated uniform(Fig. 3(a,f)) and speckle(Fig. 3(b,g)) illumination raw images at two different depths, where the in-focus information is corrupted by strong out-of-focus background. HiLo was performed with a wavelet filter size equal to one speckle-grain such that the sectioning is approximately one longitudinal speckle length. Our basic HiLo algorithm effectively suppressed background, as expected, but also manifestly suffered from residual low-frequency speckle noise (Fig 3(c,h)). In comparison, NLM HiLo benefited from the incoherent averaging of many more speckle grains, leading to largely denoised images that still preserved the intensity variations of the object itself (Fig. 3(d,i)). As a result, whereas the fine details of the pollen grain are lost in the max intensity projection (MIP) of the basic HiLo reconstruction (Fig. 3(l)), they are well preserved in the MIP of the NLM denoised reconstruction (Fig. 3(m)), as verified by comparing with the ground-truth confocal MIP (Fig 3(n)).

**Fig 3.**
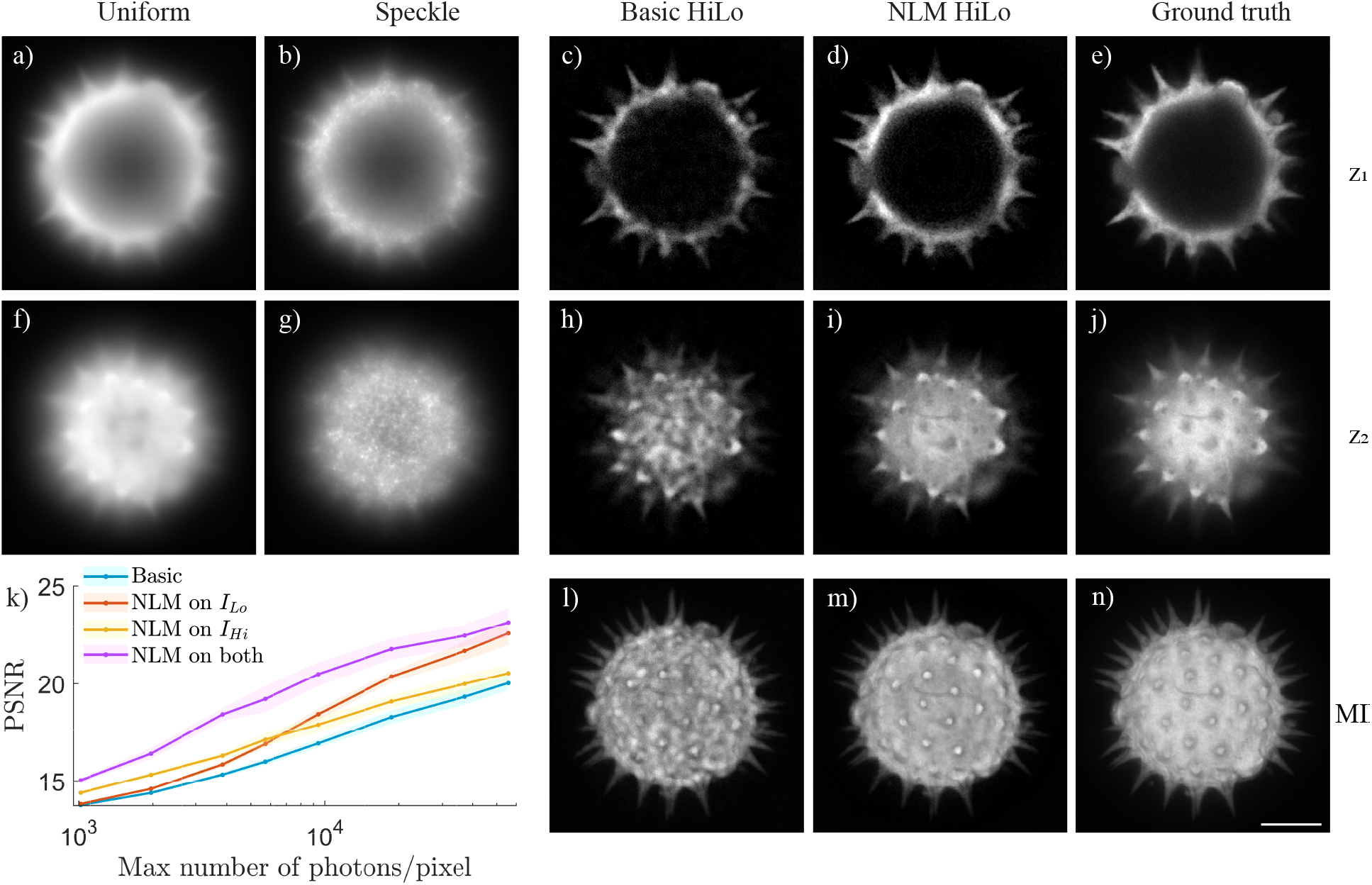
Simulation of a pollen grain stack. Images of two different depths are selected for presentation. Uniform and speckle illuminated images at *z*_1_ (a,b) and *z*_2_ (f,g). Corresponding optically-sectioned reconstructions at two depths using basic HiLo (c,h) and NLM HiLo (d,i). (e,j) Confocal images serve as a ground truth. (l-n) Maximum intensity projections (MIP) of the image stacks using basic (l) and NLM HiLo (m) compared to ground truth (n). (k) PSNR for different HiLo reconstructions as a function of signal strength. Solid line and shaded area indicate mean and standard deviation of 10 trials. Scale bar: 10 μm.

We further compared the reconstruction performance for different signal levels in Fig. 3(k), where PSNR was used to quantify the similarities between the reconstructions and ground truth. Basic HiLo was compared with NLM HiLo with denoising of *I*_Lo_ only (i), denoising of *I*_Hi_ only (ii), and denoising of both (iii). At each signal level, the PSNR of the full volume was calculated using the confocal stack as reference. The resulting PSNR was plotted as a function of the maximum number of photons per pixel in the uniform stack. At low SNR, where shot-noise variance is stronger than speckle variance, denoising of *I*_Hi_ plays a more important role than denoising of *I*_Lo_, whereas the roles are reversed at high SNR where speckle noise becomes dominant.

### 4.2 Experimental results with fixed slide

Having demonstrated NLM HiLo with simulated data, we next evaluated its performance using our multiplane imaging setup with fixed samples. We imaged a Melittobia digitata slide (Carolina Biological) with a 20*×*, 0.5 NA objective and compared the multiplane reconstructions from basic HiLo and NLM HiLo with a confocal stack taken of the same volume. The wavelet parameter was set to be σ_*w*_ = 1.5, meaning that the wavelet filter was slightly larger than a speckle grain in this case. Figures 4(a) and (b) show the raw speckle and uniform illumination images of a single plane. The high frequency images derived from *I*_*u*_ without and with NLM denoising are shown in Fig. 4(c) and (f), confirming that NLM effectively attenuates noise without compromising image resolution. The low frequency images derived from *I*_*cu*_ and denoised 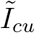 are shown in Fig. 4(d) and (g). The basic *I*_*Lo*_ was corrupted by strong speckle noise that ultimately propagated to the final HiLo image (Fig. 4(e)), overwhelming the object structure. In comparison, the speckle noise in *I*_*Lo*_ was significantly attenuated by NLM, allowing the object structure to be revealed (Fig. 4(g,h)). We also compared multiplane HiLo reconstructions in Fig. 4(i,j), where the 9-plane projections were color-coded in depth. While both methods achieved optical sectioning comparable to confocal microscopy (Fig. 4(k)), basic HiLo suffered from deleterious speckle noise that became largely attenuated with NLM HiLo.

**Fig 4.**
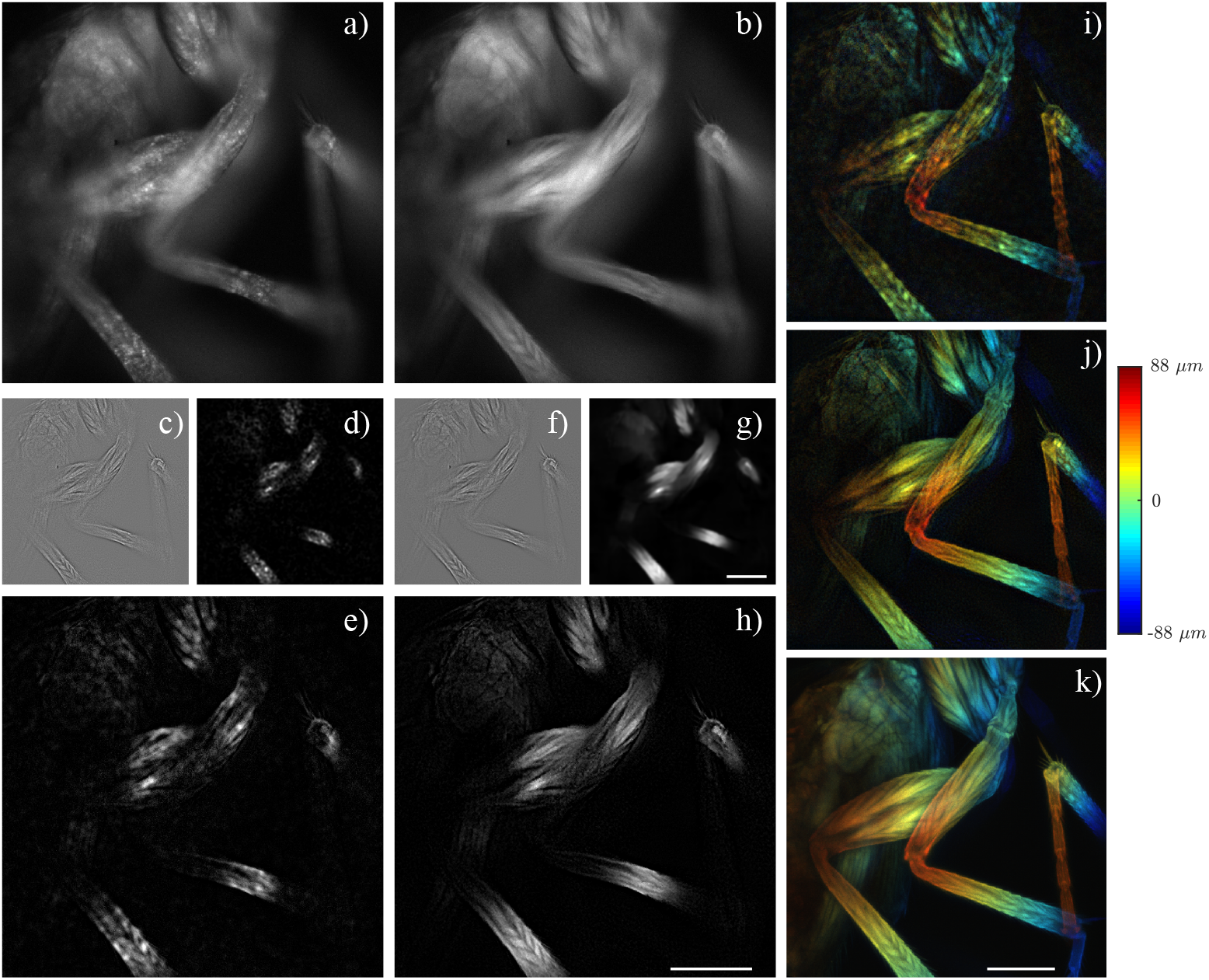
Experimental result. Multiplane HiLo imaging of a fixed slide. Widefield imaging of single plane with speckle (a) and uniform (b) illumination. Intermediate *I*_Hi_ (c), *I*_Lo_ (d) and composite *I*_HiLo_ (e) obtained with basic HiLo algorithm. Intermediate denoised *I*_Hi_ (f) and *I*_Lo_ (g) and composite *I*_HiLo_ (h) obtained with NLM HiLo. (i, j) Depthcoded projections across 9 planes from basic and NLM HiLo. (k) Depth-coded projections across a confocal stack. Scale bar: 100 μm.

### 4.3 Fast imaging of beating zebrafish heart

The zebrafish larva is a popular animal model in cardiology.^31^ Real-time contraction measurements provide important information about cardiac function and enable correlation studies with Ca^2+^ transients in the heart.^32^ Such measurements benefit from fast, volumetric and high contrast imaging, as enabled here by multiplane HiLo microscopy which provides both optical sectioning and depth information.

A beating zebrafish heart was imaged with 9-plane HiLo at a frame rate of 41 Hz (exposure time 1 ms; total recording time 17 s), using a 40*×*, 0.8 NA objective that resulted in an axial separation of 5.5 μm between planes. The results are shown in Fig. 5. Stacks of raw uniform-illumination images and NLM HiLo reconstructions are shown in the left and middle panels of Fig. 5(a). The rightmost panel of Fig. 5(a) shows a zoomed-in region comparing uniform illumination (top), basic HiLo (middle) and NLM HiLo reconstructions (bottom). NLM HiLo effectively attenuates local non-uniformities arising from speckle as well as additional noise from detection (shot and readout noise).

**Fig 5.**
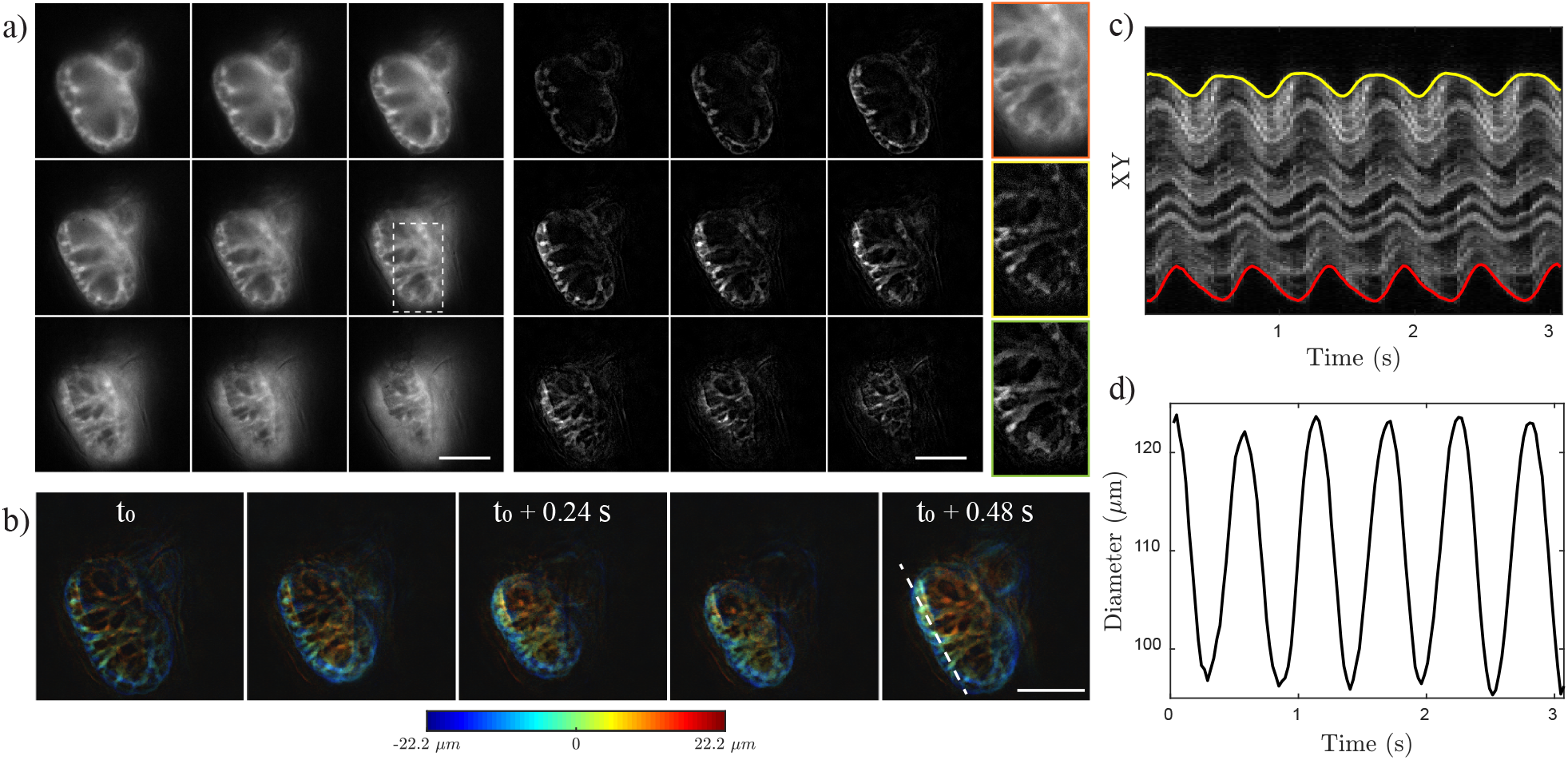
*In-vivo* imaging of beating larval zebrafish heart. (a) Multiplane stack of uniform illumination (left) and NLM HiLo reconstruction (middle). Comparison of regions outlined by rectangle for uniform illumination (top right), basic HiLo (middle right) and NLM HiLo (bottom right) images. (b) Representative snapshots of projections color-coded in depth showing cardiac contraction. (c) Kymogram obtained from line profile (dashed line in b) as a function of time. (d) Ventricle diameter as a function of time. Scale bar: 50 μm.

Representative frames showing depth color-coded projections at different times are displayed in Fig. 5 (b), where the systole and diastole are easily distinguished by virtue of the fast imaging speed. Each cardiac cycle was sampled by at least 20 images, enabling analysis of cardiac function. Here, a kymogram was determined by tracking the fluorescence signal through the ventricle (dashed line in Fig. 5(b)) and displayed in Fig. 5(c). The local ventricle diameter was calculated based on position differences of the external wall^32^and plotted as a function of time (Fig 5 (d)), illustrating different beating profiles throughout the ventricle. The heart rate was 1.8 Hz on average.

### 4.4 In-vivo calcium imaging of zebrafish brains

Finally, we illustrate the capabilities of multiplane HiLo when applied to imaging of neuronal activity in zebrafish larvae, which normally can be resolved only with confocal or two-photon. Three-plane widefield imaging of brain dynamics with optical sectioning has been demonstrated with HiLo and an electrically tunable lens (ETL).^33^Here we make use of a passive z-splitter prism to simultaneously image 9 planes with 22 μm interplane separation. Zebrafish larvae expressing nucleus-localized GCaMP6f were imaged using a 20*×*, 0.5 NA objective with 40 ms exposure time and 10 Hz frame rate. We recorded for a total duration of 73 seconds. The raw uniform and speckle illumination images were processed with NLM HiLo and the resulting video was post-processed using CaImAn^34^for calcium dynamics analysis. The results are shown in Fig. 6.

**Fig 6.**
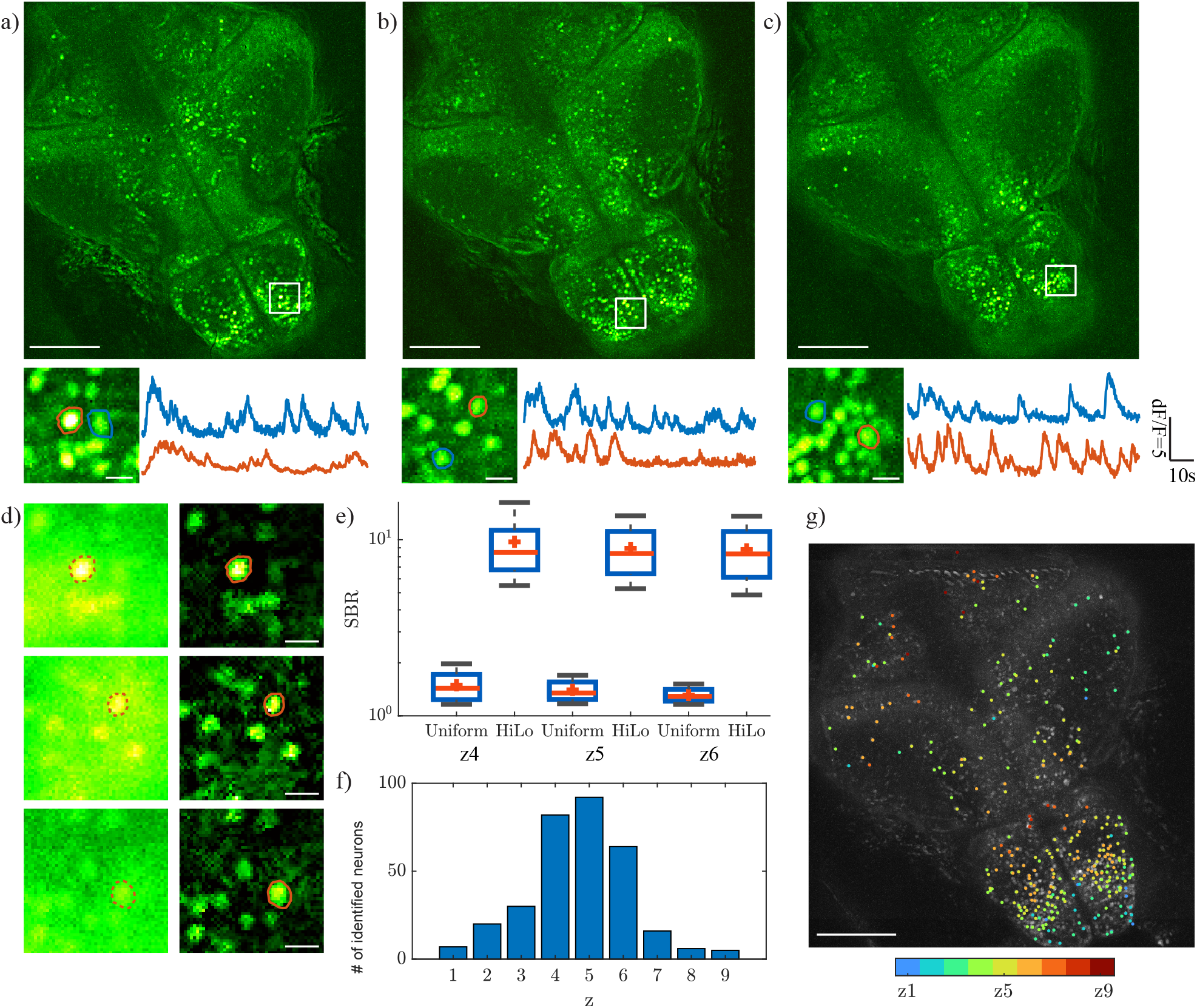
*In-vivo* multiplane calcium imaging of larval zebrafish brain. (a-c) Top: temporal MIP of HiLo frames at three different depths. Bottom: enlarged regions delimited by white rectangles showing neurons (left) and their corresponding calcium traces (right). (d) Comparison of uniform illumination (left) and HiLo (right) images for single neuron in (a-c) when active. (e) SBR for uniform illumination and HiLo images at different planes. (f) Number of identified neurons at different planes. (g) Depth color-coded neuron centroids overlaid on a HiLo MIP image. Scale bar: top panel in (a-c),(g): 100 μm; elsewhere 10μm.

The top panels in Fig. 6(a-c) display temporal MIPs obtained over the full recording duration, showing all active neurons from three central planes. The activity of individual neurons can be resolved in both space and time (Fig. 6(a-c), bottom). Single-frame widefield and HiLo images obtained from the zoomed-in rectangle regions at three depths are compared in Fig. 6(d). For the uniform illumination image, the in focus neuronal signal was easily overwhelmed by the strong background, resulting in a low signal-to-background ratio (SBR) that hampers the accuracy of neuronal segmentation. In comparison, NLM HiLo images feature lower noise and much higher contrast, by virtue of being optically sectioned. The SBR comparison of uniform illumination and HiLo images at each plane is shown in Fig. 6(e), where SBR was calculated from the ratio of the signal intensity within each neuron while active and the average background intensity within the associated circular surround region (radius 40 μm). HiLo increased SBR by nearly an order of magnitude owing to the rejection of background.

Figure 6(f) shows the total number of active neurons identified per plane over the full recording duration. Most of the neurons were identified in the three central planes (z=4-6) partly owing to the inherent distribution of neurons and partly owing to the increasing difficulty to resolve neurons at larger depths due to scattering. The identified neuron centroids, color-coded in depth, are shown in Fig. 6(g).

## 5 Discussion

The NLM denoising is different from traditional local spatial filtering in that it takes advantages of image redundancy, aiming to perform incoherent averaging between uncorrelated image patches without undermining resolution. In the case of HiLo microscopy, two images are available for the seeking of patch similarity. We have found that the similarities are more robustly found using the uniform-illumination image, which contains less noise power than the speckle-illumination image, and where the noise is more generally guided by Poisson statistics. In our case, NLM denoising is applied in the spatial domain. Other variants are also possible, combining space and frequency domain filtering^35^ or spatial-temporal information,^36^ which could potentially further improve image quality.

While the application of NLM to HiLo largely solves the problem of residual speckle noise that can arise from our original speckle-based HiLo algorithm, it does come with a drawback, namely speed. The additional time required for NLM denoising depends on the sizes of the patch search windows and of the images themselves. For example, for a 571 *×* 571 image with patch search size 31 *×* 31 pixels, the additional time required for NLM processing was 9.4 s, compared to the 0.1 s required for basic HiLo processing on a desktop computer equipped with i7-6700K CPU, 64 GB RAM. That is, NLM denoising in our case was not performed in real time. On the other hand, the physical acquisition of the raw images was fast, limited only by our camera frame rate. In particular, our two-beam illumination configuration enabled rapid switching between uniform and speckle illumination on the order of *∼*10 ms, no longer relying on the synthesis of uniform illumination by speckle averaging. This configuration also has the benefit of employing illumination patterns that are static from frame to frame. In other words, the illumination itself does not produce temporal fluctuations in the signals that could perturb measurements relying on ratios of fluorescence levels (e.g., as used in Ca^2+^ imaging).

Finally, the addition of a z-splitter prism to our setup allowed us to perform simultaneous multiplane HiLo imaging in a simple and versatile manner, using a single camera. An advantage of a z-splitter prism over alternative multiplane strategies involving the use of diffractive optical elements^19^ is that it is largely achromatic, avoiding the requirement of custom-built chromatic correctors. It is also more light efficient, which, in the case of fluorescence imaging, can be of critical importance.

In summary, we presented a multiplane HiLo microscope that provides high contrast optically-sectioned imaging with depth resolution. We also added NLM denoising to our speckle-illumination HiLo algorithm to achieve a significant reduction in residual speckle artifacts, making our imaging more accurate and robust. This was demonstrated with in-vivo imaging of both cardiac and brain activity in zebrafish larvae. Our device provides a simple solution for fast, volumetric fluorescence imaging, which can be of general interest.

## Supplemental Material

**S1 Video. *In-vivo* imaging of a beating larval zebrafish heart**. This video shows a composite video of a 9-plane color-coded z stack acquired at 41 Hz frame rate. Imaging volume: 230 *×* 230*×* 44 μm^3^.

## Disclosures

The authors declare no conflicts of interest.

## Code, Data, and Materials Availability

Associated code can be accessed at https://github.com/biomicroscopy/NLM-HiLo. Data are available from the corresponding author upon reasonable request.

## Acknowledgments

Research reported in this publication was supported by the Boston University Micro and Nano Imaging Facility and the National Institutes of Health under award Numbers R01EB029171 and S10OD024993.

## Notes

### Competing Interest Statement

The authors have declared no competing interest.

